# Tissue resident memory T cells populate the human uveal tract

**DOI:** 10.1101/2025.05.07.652385

**Authors:** Andrew D. Foers, Ian R. Reekie, Lakshanie Wickramasinghe, Amy Ward, Thomas MW. Buckley, Moustafa Attar, Soledad Aguilar-Munoa, Queen Pilapil, Maram EA. Abdalla Elsayed, Sam Pledger, Ananya Bhalla, Robert Hedley, Sarah Hill, Dylan Windell, Keith Barton, Imran Masood, Kin Sheng Lim, Mark C. Coles, Jonathan P. Sherlock, Andrew D. Dick, Sarah E. Coupland, Stephen N. Sansom, David A. Copland, Christopher D. Buckley, Srilakshmi M. Sharma, ORBIT Consortium

**Affiliations:** Kennedy Institute of Rheumatology, University of Oxford, Oxford, UK; Academic Unit of Ophthalmology, Translational Health Sciences, University of Bristol, Bristol, UK; Oxford Eye Hospital, John Radcliffe Hospital, Oxford University Hospitals NHS Trust, UK; National Institute for Health Research (NIHR) Biomedical Research Centre, Moorfields Eye Hospital NHS Foundation Trust, Moorfields, UK; Institute of Ophthalmology, University College London, London, UK; Botnar Research Centre, University of Oxford, Oxford, UK; Sir William Dunn School of Pathology, University of Oxford, Oxford, UK; Birmingham and Midland Eye Centre, City Hospital, Sandwell and West Birmingham Hospitals NHS Trust, Dudley Road, Birmingham, UK; Department of Ophthalmology, St. Thomas Hospital, London, UK; Janssen Global Services LLC, Palo Alto, California, USA; Liverpool Ocular Oncology Research Group, University of Liverpool, Liverpool, UK

**Author notes:** Co-second author (Ian R. Reekie, Lakshanie Wickramasinghe, Amy Ward).

## Abstract

The current concept is that the eye is an immune privileged site endowed with innate immune regulatory networks to maintain organ function. We now have evidence that resident T cells occupy intraocular tissues. In immune-mediated inflammatory diseases, such as psoriasis and rheumatoid arthritis, tissue resident T cells trigger disease flares in the skin and joints. This suggests resident T cells in the uvea may have similar functions in non-infectious immune-mediated uveitis, a collective term for autoinflammatory and autoimmune diseases of the uveal tract causing intraocular inflammation.

Here, we demonstrate by spectral cytometry and immunofluorescence imaging that non-inflamed uveal tissue contains multiple T cell subtypes including CD8+ CD103+ tissue resident memory T (T_RM_) cells. Using single cell RNA & T cell receptor (TCR) sequencing to profile aqueous humour cells from donors with acute, active uveitis, we identify clonally expanded T cells which are enriched for T_RM_ -associated genes. We further show that in donors with active uveitis, CD8+ CD103+ T cells persist within tissue in the uveal tract. Using bulk RNA sequencing and weighted gene co-expression network analysis (WGCNA) we show that quiescent iris tissue from donors with a history of uveitis are enriched for genes associated with T cell activation and antigen presentation. Finally, we demonstrate that T_RM_ cells persist in the anterior uvea in mice following resolution of experimental autoimmune uveoretinitis (EAU). Our results show that the human eye contains T cells both in health and during active inflammation. Our findings challenge the dogma that the eye is devoid of lymphocytes and supports the concept of resident T cell involvement in the pathogenesis of non-infectious immune-mediated uveitis and as promising targets for therapy.

**One Sentence Summary:** T cells infiltrate aqueous humour during intraocular inflammation and have capacity to migrate into uveal tissue where they remain long-lived.

## INTRODUCTION

Immune-privilege in the eye is a highly specialised function of the intraocular environment that is thought to enable local regulation of the immune response and prevent damage to vision. This is facilitated broadly by the presence of blood-ocular barriers and the presence of immunosuppressive cytokines (*1, 2*). The traditional paradigm of ocular immune-privilege was based on limited evidence for lymphocytes and lymphatics within the eye (*3*). This view is underpinned by high success rates of corneal transplantation without the need for HLA-matching or immunosuppressive therapy (*4*). Recent studies demonstrating that resident T cells occupy the normal human cornea (*5*) and naive murine anterior uvea (*6*), as well as accumulating evidence that the eye contains distict lymphatic drainage systems (*7*), challenge this view.

Tissue resident memory T (T_RM_) cells are long-lived, antigen-experienced T cells that are typically sequestered within peripheral, non-lymphoid tissues throughout the body (*8*). T_RM_ cells survey presented antigens and enable rapid responses to reinfection (*8, 9*). A consistent hallmark of their identity is expression of the cell surface markers CD69 and CD103, although this is not absolute. T_RM_ cells do not typically recirculate but can promote recruitment of central memory CD8+ T cells and other lymphocytes into tissue through local cytokine release (*9*). It is through their capacity to recognise antigen and recruit inflammatory cells into tissue that T_RM_ cells can contribute to the development of autoimmune and autoinflammatory disease. Notably, T_RM_ cells are present in white matter of human brain tissue (*10*) and are associated with chronic inflammatory and neurodegenerative brain disease (*11*). If T_RM_ cells are tolerated in the brain where, like the eye, inflammatory T cell responses are tightly controlled, it follows that they have similar functions in the eye. Evidence for this is accumulating. For example, in murine cornea, CD8+ CD103+ T_RM_ cells form following viral infection and protect against reinfection (*12*). Whereas in healthy human cornea, T_RM_ cells protect against environmental stimuli including pollen and contact lens associated damage (*5*). However, there has been limited evidence of T_RM_ cells within the human uveal tract and their contributions to uveitis.

In this study, we challenge the conventional understanding of ocular immune-privilege and suggest a mechanism for triggering relapse in ocular inflammatory disease. We describe a population of resident T cells within the normal human uveal tract. By profiling T cells within inflamed aqueous humour of infectious and non-infectious human uveitis patients, we find that clonally expanded T cells bear markers of tissue residency. We show an enrichment of T cell markers in iris tissue from quiescent uveitis donors. Finally, we demonstrate that T_RM_ cells accumulate in the anterior uvea during experimental autoimmune uveoretinitis (EAU) and persist as inflammation resolves. This work indicates that T_RM_ cells occupy uveal tissue and protect against pathogen reinfection. Meanwhile, in immune-mediated non-infectious uveitis T_RM_ cells may contribute to the development/relapse of disease and be promising therapeutic targets.

## RESULTS

### Human uveal tissue contains CD4 and CD8 T cells

To assess whether T cell profiles within the human uveal tract differ from peripheral blood during homeostasis, the CD8:CD4 T cell ratio was compared by flow cytometry. Compared to peripheral blood, uveal digests contained an elevated CD8:CD4 ratio which is consistent with the uveal tract having distinct T cell populations (Figure 1A-B). To confirm that these cells are located within tissue (and not in vessels), non-inflamed human uveal tissue sections were assessed by multiplexed immunofluorescence imaging. CD4+ and CD8+ T cells were identified within iris (Figure 1C), ciliary body (Figure 1D) and choroid (Figure 1E) and distinct from the CD34+/CD31+ vasculature.

**Figure 1.**
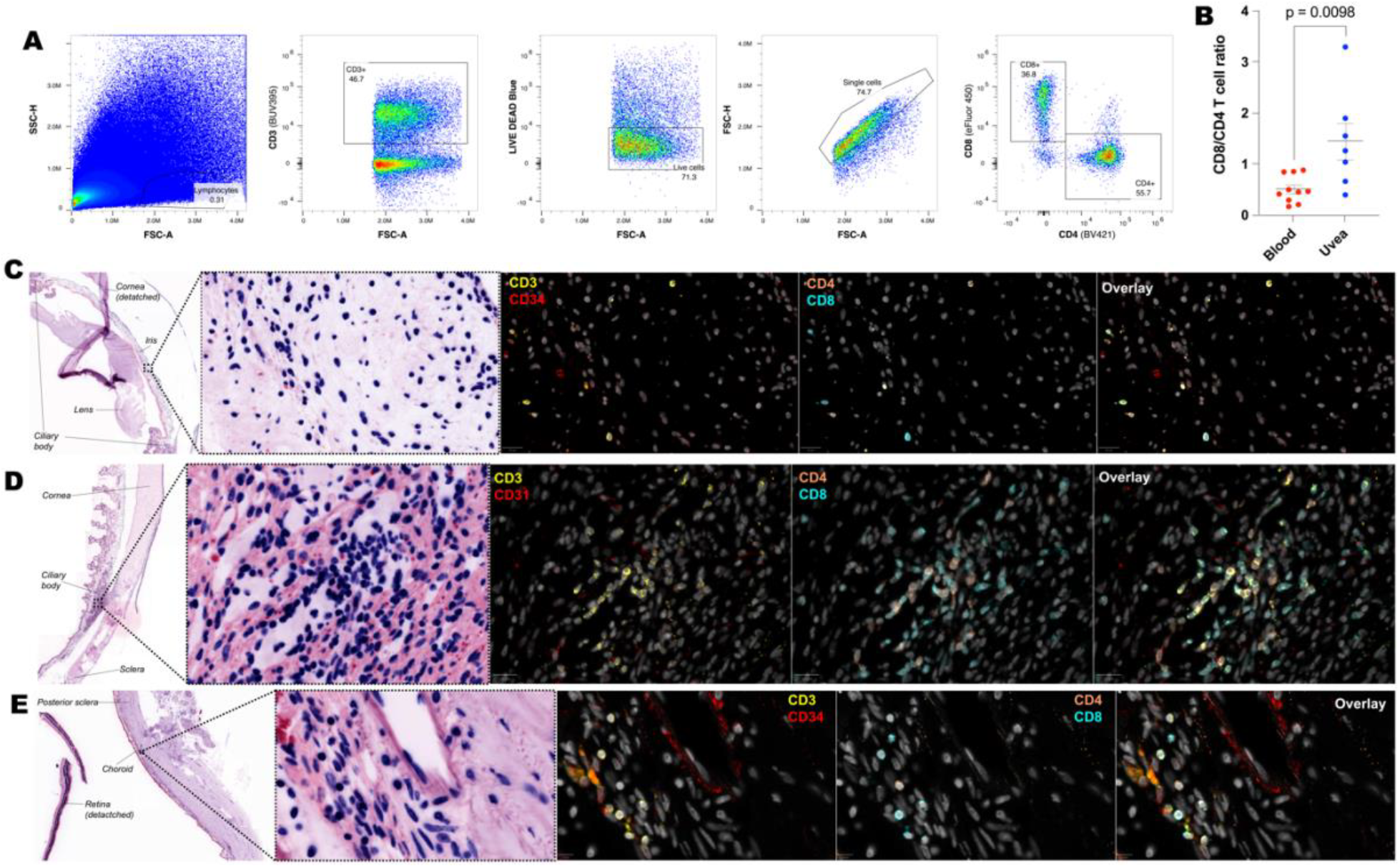
Uveal tissue contains CD4 and CD8 T cells. **(A)** Gating strategy for selecting live CD4 and CD8 T cells. **(B**) The ratio of CD8:CD4 T cells is increased in uveal digests compared to blood. Immunofluorescence confirmation of CD4 and CD8 T cells within **(C)** iris, **(D)** ciliary body and **(E)** choroid tissue and distinct from CD31+/CD34+ blood vessels.

### T_RM_ cells populate the human uvea

To further phenotype T cells within the human uveal tract, CD3+ cells were enriched from the uvea of 4 post-mortem human donors without overt ocular disease (Supplementary Table 1), and examined using a 25-marker T cell panel for analysis by spectral flow cytometry (Figure 2A). In total 17 T-cell subpopulations were defined comprising 7 clusters of CD8 T-cells, 5 clusters of CD4 T cells and 5 clusters of double-negative γ*δ* T cells (Figure 2B-C; Supplementary Figure 1). Proportions of each cell type were comparable across the 4 donors (Figure 2D). The CD8 T-cells comprised: (i) two clusters of TCR-Va7.2 CD161hi MAIT cells (C4, C5), one of which expressed the T_RM_ marker CD103 (C5), (ii) two clusters of CD69hi cells which expressed the T_RM_ markers CD49a (C7) and CD103 (C3), (iii) CD25+ memory (or regulatory) cells (C6), CD27/28neg CD45RAhi terminally differentiated effector memory (T_EMRA_) T cells (C13) and CCR4/6hi central memory cells (C9). The CD4 T cells comprised: (i) two CD25hi regulatory cell clusters (C1, C2) one of which expressed CD103 (C2), (ii) CD45ROneg naïve cells (C10), CCR7/10hi Th cells (C11) and CCR6hi Th17 effector memory cells (C8). Finally, the double-negative γ*δ* T cells comprised (i) two clusters of Vγ9/V*δ*2 cells with CD27hi naïve (C16) and CCR10 Th (C15) like phenotypes and (iii) three clusters of V*δ*1 cells with CD45RAhi naïve (C17) CCR10 Th (C12) and CCR6/CXCR3/CCR7/IL-23Rhi cells (C14). Notably, high expression of IL-23R in C14 suggests this might be a resident γ*δ* T cell population pathogenically linked to spondyloarthropathy (*13*) and anterior uveitis (*6, 14*).

**Figure 2.**
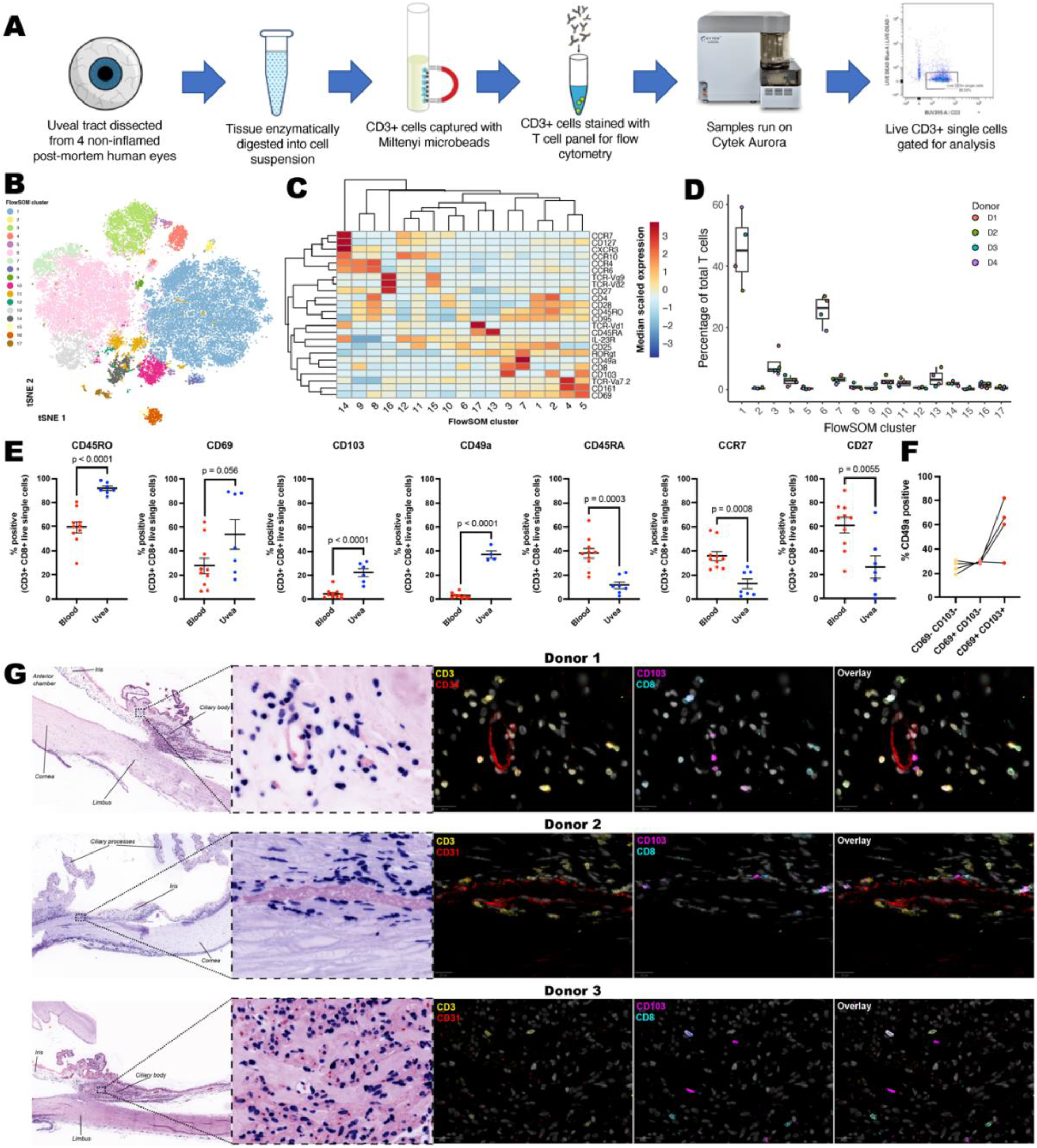
T_RM_ cells populate non-inflamed human uveal tissue. **(A**) Experimental overview for spectral cytometry. **(B**) tSNE plot of individual cells coloured according to FlowSOM cluster. **(C**) Heatmap depicting expression of T cell genes in FlowSOM clusters. **(D**) Boxplots depicting composition of the T cell compartment across the 4 donors. **(E)** Comparison of T_RM_ cell marker genes between live uveal and blood CD3+ CD8+ cells. **(F)** CD49a expression within CD8+ CD69 and CD103 T cell subsets. **(G)** Confirmation of CD3+ CD8+ CD103+ cells within histologically normal non-inflamed uveal tissue by immunofluorescence imaging.

To confirm that cells positive for T_RM_ markers are *bona fide* tissue resident cells, and not contaminants from the vasculature, uveal and peripheral blood CD3+ CD8+ cells were compared by flow cytometry for residency and circulatory markers (Figure 2E). Compared to blood, CD3+ CD8+ cells in uveal digests were enriched for CD45RO, CD69, CD103 and CD49a, and depleted for CD45RA, CCR7 and CD27 supporting the presence of canonical T_RM_ cells within normal human uveal tissue. CD8+ CD69+ CD103+ uveal T_RM_ cells varied in their expression of CD49a (Figure 2F), indicating that uveal T_RM_ cells have capacity to produce both IFNγ and IL-17 (*15*).

To further confirm the presence of T_RM_ cells, non-inflamed human anterior uveal tissue sections were assessed by immunofluorescence imaging (Figure 2G). CD3+ CD8+ CD103+ cells were located adjacent to vasculature (Donors 1 & 2), including at the iris root (Donor 1) and deep within ciliary muscle (Donor 3). Numerous CD4+ and CD8+ T cells were also apparent that did not express CD103, and may relate to other T cell subsets described above, and previously reported CD8+ CD103-T_RM_ cells (*9*).

### Expanded T cell clones within the aqueous humour of uveitis donors are enriched for T_RM_ genes

To profile cells within the ocular fluid of donors with uveitis, aqueous humour was collected from 11 donors with active uveitis and cells analysed by scRNAseq with V(D)J sequencing (Figure 3A-B; Supplementary Table 2). Broad cell types were initially annotated (Figure 3C) with T cells being the predominant cell population in 9 of 11 donors (Supplementary Figure 2). The 2 donors in which T cells were not the predominant cell type were diagnosed with HLA-B27+ acute anterior uveitis and their aqueous humour cells predominately comprised monocytes. This result is consistent with a previous report showing that myeloid cells predominate in the aqueous humour of HLA-B27+ acute anterior uveitis (AAU) patients (*16*).

**Figure 3.**
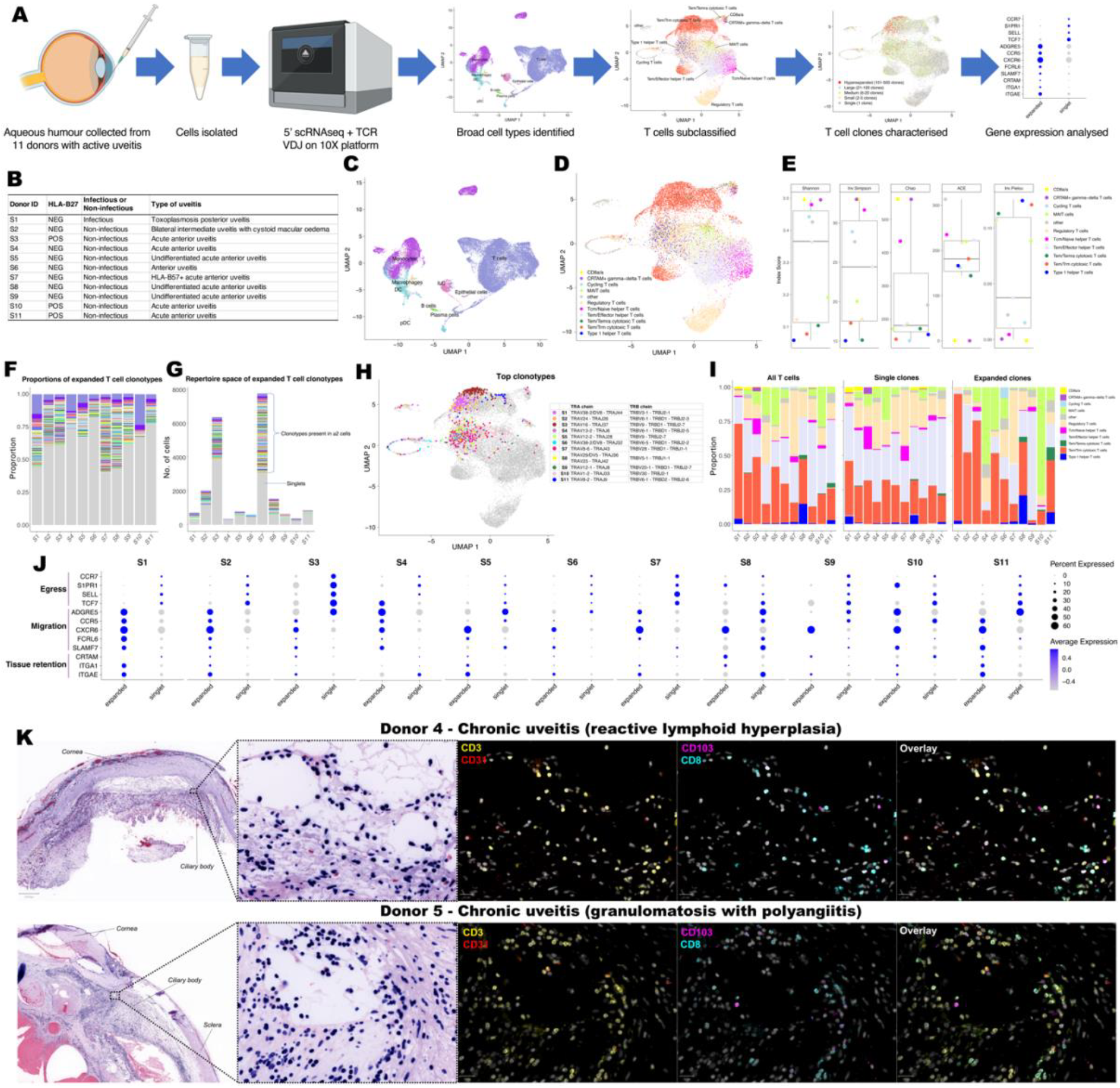
T_RM_ cells populate human aqueous humour and uveal tissue during active uveitis. **(A)** Experimental overview for scRNA sequencing and T cell receptor profiling of cells in aqueous humour during uveitis. **(B)** Uveitis diagnosis and HLA-B27 status of aqueous humour donors. UMAP plots of **(C)** all cells with broad cell types annotated and **D**) T cell subtypes following subsclustering of T cells only. **(E)** Analysis to different diversity metrics to assess diversity of T cell receptor repertoire across T cell subtypes. **(F)** Proportions and **(G**) repertoire space of expanded T cell receptor clonotypes. **(H)** For each donor, cells expressing the most abundant clonotype are highlighted. Corresponding TRA and TRB chains of abundant clones are specified. **(I)** Comparison of T cell subtype proportions between all T cells, single clones and expanded (present in ≥2 cells) clones. **(J)** Comparison of egress, migration, and tissue retention markers between expanded and non-expanded T cells. **(K)** Immunofluorescence imaging of CD3, CD8 and CD103 cells in enucleated eyes from donors with advanced uveitis donors alongside the vascular marker CD31.

To further characterise T cell populations, cells broadly classified as a T cell that also expressed a T cell receptor (TCR) were subclustered. 10 different T cell subtypes were identified, with T_EM_/T_RM_ cytotoxic T cells the most prevalent (Figure 3D). Since clonally expanded T cells within the synovial fluid of patients with spondyloarthropathy contain a T_RM_ signature (*17*), we hypothesised that this may also occur in the aqueous humour of patients with uveitis. To this end, paired TCR*α* and TCR*β* CDR3 amino acid sequences (clonotypes) were annotated for each cell. T_EM_/T_RM_ cytotoxic T cells had low levels of TCR diversity compared to other cell types (Figure 3E), consistent with involvement of these cells in antigen recognition. Strikingly expanded clones (clonotypes present in ≥2 cells) made up 17-60% of the repertoire space for each donor (Figure F-G). To investigate the T cell subtype and TCR*α*/TCR*β* chains driving antigenic responses, the most abundant clonotypes in each donor were further investigated (Figure 3H). For donors, S1, S2, S3, S5, S11 the most abundant clonotype comprised T_EM_/T_RM_ cytotoxic T cells; for donors S4, S6, S10 the most abundant clonotype were MAIT cells; for donors S7, S8 the most abundant clonotype were regulatory T cells; and for S9 the most abundant clonotype were T_EM_/Effector helper T cells. Although variable across the individual donors, T_EM_/T_RM_ cytotoxic T cells were more likely to harbour expanded clonotypes compared to singlets (T cells with unique clonotypes) (Figure 3I). There was also a notable MAIT cell clonal expansion in donors S4 (HLA-B27-AAU) and S10 (HLA-B27+ AAU) (Figure 3I). Overall, cells harbouring an expanded clonotype were enriched for genes associated with cell migration and tissue retention and depleted of egress genes, compared to singlets (Figure 3J).

We next investigated if T_RM_ cells are present within uveal tissue during active uveitis. For this, enucleated eyes from 2 donors with end-stage uveitis (Supplementary Table 3) were analysed by immunofluorescence imaging. Both donors displayed abundant tissue infiltrating CD3+ cells including CD8+ CD103+ cells in the anterior uvea (Figure 3K). Notably, CD8+ CD103+ cells were also present in the retina in chronic end-stage uveitis, whereas in control tissue with no history of uveitis, no T cells were present within the retina (Supplementary Figure 3).

### Iris tissue contains T cell and antigen presenting gene signatures during quiescent uveitis

We next investigated if T_RM_ cells persist in iris tissue during uveitis remission. To this end, iris tissue was collected from 13 donors, 6 with glaucoma secondary to a history of uveitis (now quiescent following treatment) and 7 glaucoma donors without uveitis (Figure 4A; Supplementary Table 4). Iris tissue was processed for bulk RNA sequencing and weighted gene co-expression network analysis (WGCNA) was performed. Genes with similar expression were grouped into modules and module eigengenes (MEs) calculated (Supplementary Figure 4A-B; Supplementary Table 5). Based on an alpha of 0.05, three modules positively and two modules negatively correlated with donors that had a history of uveitis (Figure 4B; Supplementary Figure 4C). Gene ontology (GO) analysis was applied to characterise the pathways associated for genes within modules of interest (Figures 4C-E; Supplementary Table 6). GO analysis identified the black module as enriched for genes associated with T cell signalling. Consistent with long-lived T cells persisting in uveal tissue following resolution of uveitis, genes in the black module were enriched for T cell associated genes and responsiveness to interleukin 7. This included CD8A/CD8B and CCL5 (RANTES) which were amongst the genes most strongly associated with the black module (Figure 4F). The brown and green modules were respectively enriched for genes associated with antigen presentation and angiogenesis, possibly reflecting the previously inflamed state of the tissue.

**Figure 4.**
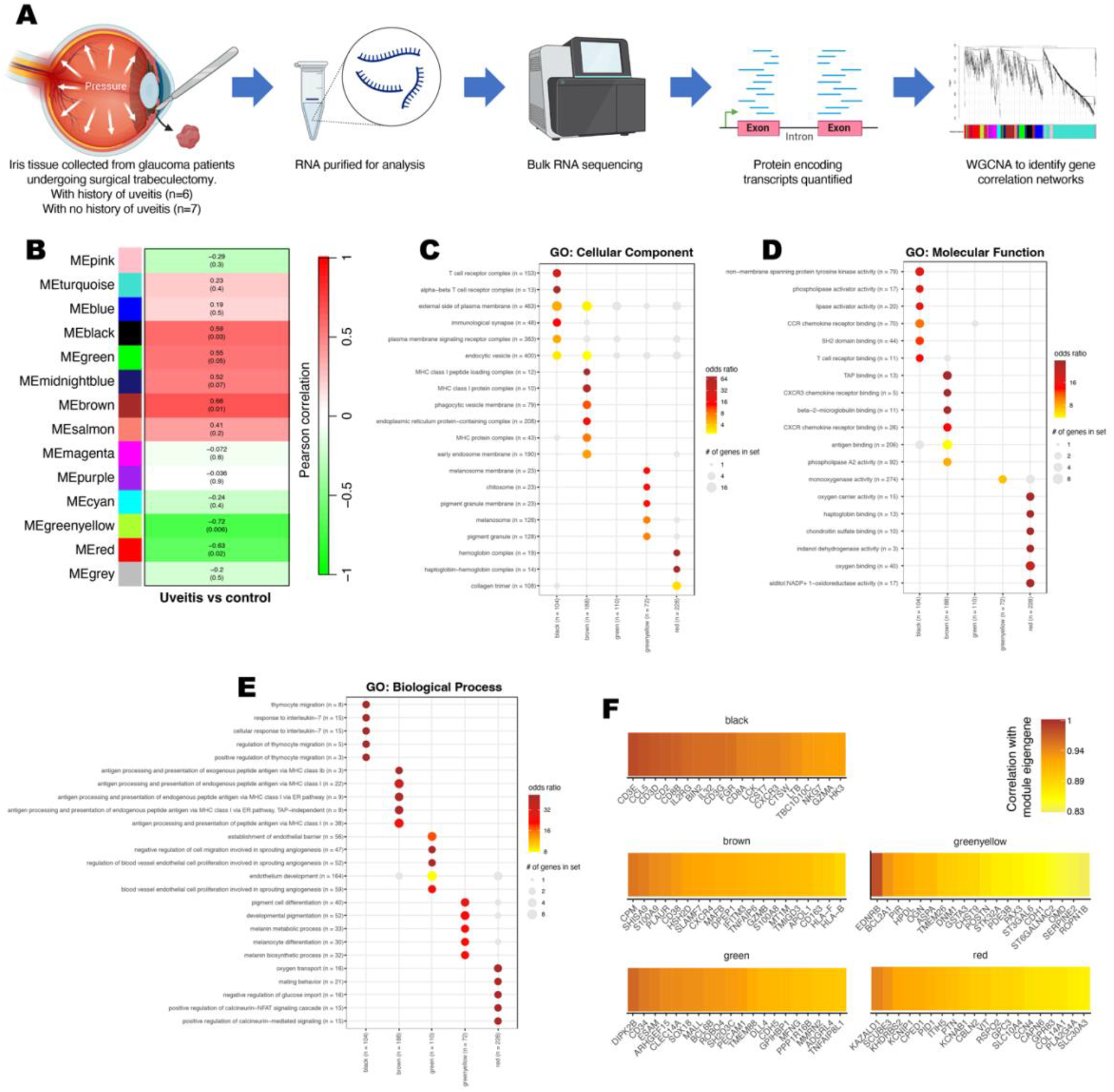
Previously inflamed iris tissue is enriched for genes associated with activated T cells and antigen presentation. **(A)** Methodological overview to compare gene expression profiles within iris tissue recovered during glaucoma surgery from donors with and without a history of uveitis. **(B**) Weighted gene co-expression network analysis (WGCNA) of correlations with module eigengenes (MEs) and uveitis. Pearson correlations are shown with corresponding P value in brackets. Gene ontology (GO) analysis of **(C)** cellular components, **(D)** molecular functions and **(E)** biological process associated with module genes. Number of genes within modules and GO terms are specified. GO terms with BH adjusted P values <0.05 are coloured. **(F)** Top 20 module eigengene - gene correlations.

To investigate whether the individual modules might encompass specific cell types, module genes were assessed against an ocular scRNAseq dataset (*18*). The black and green modules, which were positively correlated with uveitis donors, were respectively enriched for genes from activated T cells and endothelial cells (Table 1). Conversely, the red module, which was negatively correlated with uveitis donors, was associated with WIF1 high fibroblasts. As WIF1 is a negative regulator of WNT signalling – an important pathway in development of memory T cells – a depletion of WIF1 fibroblast in iris tissue of uveitis is consistent with elevated WNT signalling in uveitis and subsequent promotion of T_RM_ cell development (*19*).

**Table 1.** Module genes associated with eye cell types.

| Module | Gene set | P value<br>(BH-adjusted) | Odds<br>ratio | Module genes in gene set |
| --- | --- | --- | --- | --- |
| black | Iris ciliary body activated t cells<br>(n = 128) | 0.00139 | 12.2 | SPN, ADAM8, SLAMF6, IL2RB,<br>PRKCB, S1PR4, SLAMF1 |
| black | Choroid sclera activated t cells<br>(n = 119) | 0.026 | 8.53 | SIT1, FCRL6, FASLG, SLAMF1,<br>SPN |
| brown | Iris ciliary body fibroblasts<br>(n = 102) | 0.286 | 2.86 | SOD2, STEAP4, BMP1, IGFBP4,<br>KLF2, MT1A, GEM |
| green | Choroid sclera choroid endothelial cells<br>(n = 141) | 0.0594 | 8.85 | NRARP, BEX5, BCL6B, CA4 |
| green | Iris ciliary body ciliary body endothelial cells<br>(n = 356) | 0.184 | 5.44 | BCL6B, CISH, SLC52A3, IL15RA |
| greenyellow | Cornea melanocytes<br>(n = 261) | 0.422 | 4.58 | DIPK1C, STXBP1, CDH1 |
| red | Iris ciliary body WIF1 high fibroblasts<br>(n = 123) | 0.026 | 4.25 | KCNAB1, GPR37, LSAMP, H1-0,<br>RSPO2, CDH11, CTSC, DPYSL3,<br>ECRG4, LUM |

### T_RM_ cells persist in the anterior uvea of EAU mice as disease resolves

To test the hypothesis that T_RM_ cells remain in the eye as inflammation resolves, we employed the adoptive transfer EAU murine model of uveitis and assessed T cell infiltrate over the course of disease. In this non-infectious model, a predominately posterior uveitis is initiated in naïve mice following the adoptive transfer of leukocytes from donor mice immunised with the uvetiogenic peptide retinol binding protein-3 (RBP-3)_629-643_ (Figure 5A)(*20*). The peak of disease occurs at day 12, with inflammation resolving by day 49. Consistent with a posterior foci, fundus and optical coherence tomography (OCT) images of the retina reveal cellular infiltrate into the posterior chamber and destruction of retinal tissue (Supplementary Figure 5).

**Figure 5.**
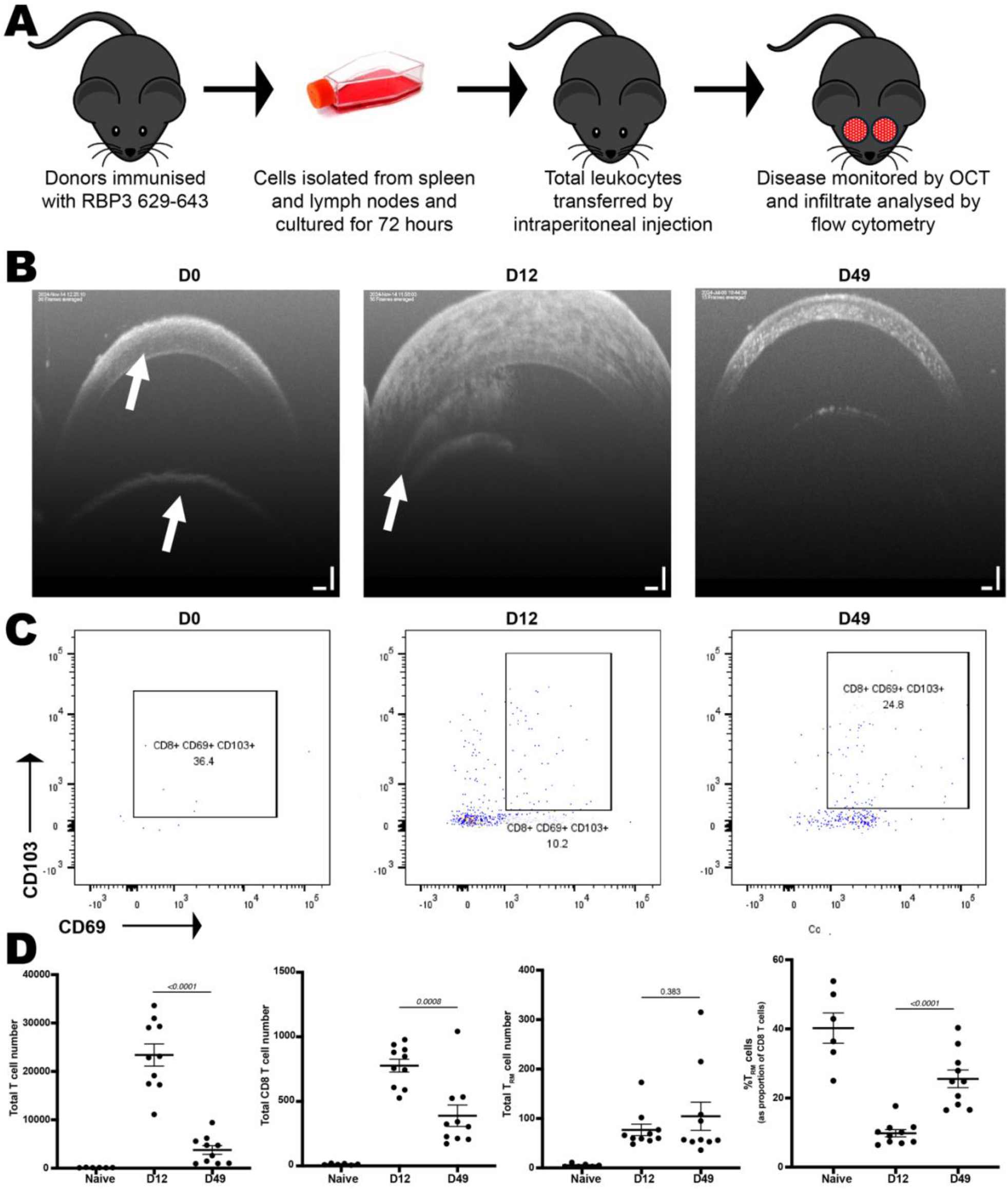
T_RM_ cells persist in the anterior uvea in post-EAU mice. **(A)** Schematic overview of adoptive transfer murine model of uveitis. **(B)** Representative OCT images of the anterior uvea with arrows to illustrate cornea and lens (D0) and angle (D12). **(C)** Representative flow cytometry plots showing CD69+ CD103+ T_RM_ cells as proportion of total CD45+ CD3+ CD8+ T cells at indicated time points. Total anterior uveal numbers of **(D)** CD45+ CD3+ T cells, CD45+ CD3+ CD8+ T cells, and CD45+ CD3+ CD8+ CD69+ CD103+ T_RM_ cells were quantified in murine donors by flow cytometry at indicated time points.

Despite being a predominately posterior model, the anterior uvea demonstrates substantial cellular infiltrate at day 12 which resolves by day 49 (Figure 5B). To investigate if T_RM_ cell involvement, mice were culled at days 0 (naïve mice; n = 6), 12 (peak disease; n = 10) and day 49 (resolved; n = 10) and anterior uveal T cells were collected, following perfusion, for flow cytometric analysis (Figure 5C). In contrast to total CD3+ T cell and CD8 T cell numbers, which decreased significantly from day 12 to day 49, T_RM_ (CD45+ CD8+ CD69+ CD103+) cells remained elevated (Figure 5D). At day 49 the CD8 T cell compartment consisted of 16-40% T_RM_ cells, despite normal appearance of the anterior chamber (Figure 5B). Strikingly, high proportions of T_RM_ cells were also apparent in naïve mice and aligns with above observations of long lived T_RM_ populations in adult human uvea without overt ocular disease.

## DISCUSSION

In this study we demonstrate that uveal tissue contains T_RM_ cells. These findings, along with the recent description of T cells in the human cornea (*5, 12*), challenge the classical view that normal uveal tissue is devoid of lymphocytes. Our study adds to accumulating evidence that structures in the eye experience different levels of ‘immune-privilege’ (*21-23*). To protect vision, the retina must maintain a high degree of immunotolerance whereas the uveal tract, cornea and sclera may tolerate and possibly benefit from the presence of some types of lymphocytes. Greater permeability of the blood aqueous than the blood retinal barrier, and ‘leakiness’ of choroidal blood vessels, are likely to facilitate these differences (*24*). This is also representative of other immune-privileged sites. For example, T_RM_ cells accumulate in the choroid plexus in the brain which has a less restrictive blood barrier than parenchymal tissue (*25*). Likewise, in the testes, normal seminiferous epithelium (the site of spermatogenesis) is considered to be totally absent of lymphocytes, whereas T_RM_ cells are present in other testicular sites (*26*). A breakdown in these barriers such as during local infection, trauma or severe inflammation would permit lymphocyte infiltration and facilitate formation of long-lived T_RM_ cells. This is evident in the eyes with T cell infiltrate in the retina of a human eye with severe chronic uveitis, but not in the retina of an eye without uveitis (Supplementary Figure 3). Similarly, there is an accumulation of T_RM_ cells in the retina in mice during chronic experimental autoimmune uveoretinitis (*27*).

Targeting T_RM_ cells may offer new approaches for preventing uveitis. T_RM_ cells at other sites are implicated in the pathogenesis of autoimmune and autoinflammatory diseases and may contribute to disease relapse. Recently, synovial CD8 T_RM_ cells were demonstrated to initiate relapse in murine models of inflammatory arthritis and depleting joint specific T_RM_ cells during remission attenuated disease recurrence (*28*). In humans, T_RM_ targeting therapy has shown promise in psoriasis and other skin conditions that are characterised by recurrent lesions in same locations (*29*). T_RM_ cell-depleting therapy potentially has value in other relapsing T-cell mediated, uveitides, such as spondyloarthropathy-associated anterior uveitis and Bechet’s uveitis.

T_RM_ cell depletion may likewise prevent certain instances of immune-checkpoint inhibitor (ICI) uveitis. Most patients on ICI therapy experience an immune-related adverse event (irAE) that is reminiscent of autoimmune disease (*30*). Notably, there is a higher incidence of irAEs in people with pre-existing autoimmune or autoinflammatory conditions where they can manifest as flare at previously inflamed sites (*30-32*). ICI-uveitis occurs in about 1% of patients on ICIs for unrelated malignancy (*33*). ICIs block T cell inhibitory proteins, including CTLA-4 and PD-1 which are highly expressed on T_RM_ cells (*34*). ICI-induced T_RM_ activation mechanistically underpins some irAEs. For example, CD8+ T_RM_ cells are key effector cells in ICI-colitis (*35, 36*) and are implicated in ICI-dermatitis (*37*). This suggests that activation of T_RM_ cells can cause ICI-uveitis – notwithstanding other ICI-induced mechanisms such as suppression of Tregs, and cross-reactivity of tumour-targeting activated T cells (*38*).

The presence of clonally expanded T_RM_ cells within aqueous humour in active uveitis informs our understanding of TCR-antigen interactions that promote uveitis. Our observations align with reports in other settings. For example, T_RM_ cells accumulate in both synovial fluid (*17*) and joint tissue (*28*) in arthritis. Likewise, during chronic neuroinflammatory and neurodegenerative diseases T_RM_ cells accumulate in cerebral spinal fluid (*11*) and central nervous system tissue (*39*). Given multiple expanded T cell clonotypes within all aqueous humour donors, it is possible that following an initial trigger that facilitates T cell influx, inflammation-associated autoantigens contribute to the local activation of infiltrating T cells. Following activation these cells upregulate T_RM_ genes which facilitates their migration into tissue. There, they remain long-lived and are positioned to promote subsequent episodes of uveitis following an inflammatory trigger (e.g., microtrauma).

Our data indicates that in addition to T_RM_ cells, other T cell subtypes reside in the uveal tissue. Both flow cytometry and immunofluorescence revealed, numerous CD3+ CD103-cells within ocular tissue indicating that additional T cells subtypes reside in the uveal tract. Of particular interest are the TCR-Vd1+ IL-23R+ population identified by flow cytometry. This T cell population resides in entheses and are linked to the development of spondyloarthropathy (*13*), and possibly spondyloarthropathy-associated uveitis (*6, 40*). There are several structures within the eye which might be considered analogous to an enthesis, such as the scleral spur or the iris root, that might harbour this T cell population. Unfortunately, our attempts to confirm the presence of these cells by immunofluorescence imaging and spatially locate them to ocular entheses were thwarted by the scarcity of these cells in non-inflamed human tissue combined with challenges in staining for IL-23R and γ*δ* T cells. We detected MAIT cells in non-inflamed control tissue as well in uveitic aqueous humour where they were highly expanded in 3 donors, each with anterior uveitis. Further work is needed to confirm if these cells have a role in reducing inflammation, as described in experimental autoimmune uveitis (*41*).

The availability of healthy ocular tissues presents a challenge for human eye research. Our study may be limited in that some of our control tissue may not truly represent normal eyes. In particular, non-uveitic tissue used for immunofluorescence was obtained from enucleated eyes containing posterior ocular melanoma, or from post-mortem donors (Supplementary Table 3). While the uveal tract and surrounding structures in these donors appear histologically normal, they may not fully reflect healthy tissue.

Until recently, it was thought that T cells were only recruited to tissues following infection or damage and then underwent egress or apoptosis following resolution of inflammation (*8*). It is now evident that long-lived resident T cells populate most tissues throughout the body and have important physiological functions. We suggest a contributory role for T_RM_ cells in the pathophysiology of immune-mediated inflammatory diseases including in the eye. Future studies are warranted to clarify their involvement in uveitis and if targeting them therapeutically improve control of some uveitides and their associated systemic diseases.

## MATERIALS AND METHODS

All human samples used in this work was approved under Research Ethics Committee number: 07/H0706/81 in the United Kingdom. Mouse work was conducted in concordance with the United Kingdom Home Office licence (PPL PP9783504) and approved by the University of Bristol Animal Welfare & Ethical Review Group. The study also complied with the Association for Research in Vision and Ophthalmology (ARVO) Statement for Use of Animals in Ophthalmic and Vision research.

### Enrichment of ocular CD3+ cells for flow cytometry

Post-mortem human eyes were obtained from NHS Blood and Transplant (ethics number: 07/H0706/81) following removal of corneas (Supplementary Table 1). For each donor, the uveal tract (including retinal pigment epithelium) was dissected, placed in 20 ml RPMI containing 1 mg/ml Collagenase II, 0.5 mg/ml DNase I and minced with scissors. To digest tissue, tubes were placed in a ThermoMixer set to 500 RPM and 37°C for 1 hour. To aid the breakdown of tissue, samples were mixed by pipetting up and down approximately thirty times every 20 minutes. The digestion was stopped by filtering the sample through a 70 µm cell strainer followed by addition of 20 ml of RPMI containing 20% FBS. Cells were spun down at 500x*g*; 5mins, washed twice in PBS and pellets resuspended in 320 µl of PBS, 0.5% BSA, 2 mM EDTA + 80 µl of CD3 MicroBeads (Miltenyi Biotec; 130-050-101). CD3 cells were positively selected for following manufacturer’s protocol and using LS columns (Miltenyi Biotec; 130-042-401) and a QuadroMACS™ Separator (Miltenyi Biotec; 130-042-401). To obtain a more stringent enrichment, each sample was passed through two LS columns. CD3 enriched cells were stained with 1:300 live-dead fixable blue (ThermoFisher; L34961) or Zombie NIR (Biolegend; 423105) in PBS for 20 mins, washed in PBS, and resuspended in 49 µl PBS, 0.5% BSA, 2 mM EDTA, and 1 µl Fc Block (Biolegend; 422301) and left for 10 mins at RT. All antibodies, except RORgt, were then directly added to cell suspensions and stained for 30 mins in dark at RT. Cells were then pelleted (500x*g*; 5 mins) and prepared for RORgt staining using the True-Nuclear™ Transcription Factor Buffer Set (Biolegend; 424401). For RORgt staining, cells were fixed in 100 µl Fix Concentrate for 45 mins at RT. Cells were pelleted, washed and permeabilised in 100 µl of the Perm Buffer and then RORgt antibody directly added to Perm Buffer and cells stained for 30 mins in the dark at RT. Finally, cells were pelleted, washed and resuspended in 3% FBS/PBS and stored in the dark at 4°C before analysing on a Cytek Aurora equipped with 5 lasers. See Supplementary Table 7 for antibody details.

### Preparation of human blood cells for flow cytometry

Peripheral blood was collected from donors in EDTA tubes and centrifuged at 300xg for 10 mins. Plasma was removed and remaining fluid diluted 1:10 with ACK lysis buffer and gently rocked for 20 mins at RT to lyse red blood cells. Leukocytes were pelleted and resuspended in 10% DMSO/90% FBS and frozen in a Mr Frosty at -80°C before long term storage in liquid nitrogen. To stain cells by flow cytometry, blood cells were thawed at 37°C, transferred to RPMI containing 10% FBS, pelleted, resuspended in complete RPMI and rested for 1 hour at 37°C. Cells were then pelleted and resuspended in 1 ml RPMI containing 1 mg/ml Collagenase II, 0.5 mg/ml DNase I and placed in a ThermoMixer set to 500 RPM and 37°C for 20 mins. To aid the digestion, samples were mixed by pipetting after 10 minutes. The digestion was stopped by adding 1 ml of 20% FBS/PBS and cells were centrifuged and pelleted resuspended and washed in PBS. To identify dead cells, cells were stained with 1:300 Zombie NIR/PBS for 20 mins in the dark. Cells were. then washed and resuspended in 49 µl PBS, 0.5% BSA, 2 mM EDTA, and 1 µl Fc Block (Biolegend; 422301) and left for 10 mins at RT. Cells were stained with antibodies (Supplementary Table 7) for 30 mins in the dark at RT. Following staining, cells were centrifuged (500xg; 5 mins) and pellets resuspended in 100 µl of fixation buffer (Biolegend; 420801) and fixed for 30 mins in the dark at RT. Cell were then pelleted by centrifugation (500xg; 5 mins), washed once and resuspended in 3% FBS/PBS and stored in the dark at 4°C before analysing on a Cytek Aurora (5 laser) spectral flow cytometer. Following acquisition, data was unmixed in SpectroFlo software (Cytek) and exported as FCS files.

### Analysis of human flow cytometry data

FCS files were imported into FlowJo (BD). Lymphocyte sized events were gated by FSC/SSC profiles followed by gating CD3+ live-dead negative, single cells which were sent for analysis; unmixed mean fluorescence intensity (MFI) values are included in Supplementary Table 8. For high parameter analysis, dimensionality reduction was performed in FlowJo using the tSNE function, and cells grouped into clusters with the FlowSOM plugin (*42*). tSNE values, FlowSOM clusters and MFI values were imported into R (v4.3.1) for visualisation. tSNE plots were created with ggplot2 (v3.4.3) and median scaled MFI values used to create the heatmap using the pheatmap package (v1.0.12).

### Immunofluorescence imaging

Formalin-fixed paraffin embedded (FFPE) tissue slides from 5 donors (Supplementary Table 3) were deparaffinised, rehydrated and prepared for imaging on the GE Cell Dive multiplex platform as previously described (*43*). Slides were antigen retrieved in a citrate antigen unmasking solution (Vector Labs; H3300) in a pressure cooker at 95°C for 20 mins before being washed in PBS and blocked in 10% donkey serum/3% BSA/PBS. Following blocking, slides were washed in PBS and stained with DAPI (ThermoFisher, D3571) for 15 min and washed again. Slides were cover slipped in mounting media (50% glycerol (Sigma; G5516) + 4% propyl gallate (Sigma; 2370)). Slides were first imaged at 10X to acquire a scan plan of regions of interest followed by imaging at 20X to establish background autofluorescence and generate virtual H&E images. Coverslips were then removed by submerging slides in PBS and tissue was Fc blocked in 1:200 FcR blocking reagent (Miltenyi Biotec; 130-059-901) in 3% BSA/PBS. All antibody cocktails were diluted in 1:200 FcR blocking reagent in 3% BSA/PBS and tissue was stained overnight at 4°C. Antibody details are specified in Supplementary Table 9. Primary antibodies with corresponding secondary antibodies were used in the first round of imaging only. After a round of image acquisition, fluorophores were bleached three times in 0.1 M NaHCO_3_ (Sigma; S6297), 3% H_2_O_2_ (Merck; 216763), pH 11.2 for 15 each with a 1 min PBS wash in-between each bleaching round. Slides were restained with DAPI for 2 mins, washed and coverslipped to acquire background fluorescence. Coverslips were then removed and tissue stained with directly conjugated secondary antibodies. After washing 3 times in PBS, slides were coverslipped and images acquired. Multiple imaging rounds were overlaid and multiplexed images imported into QuPath for visualisation.

### Single cell RNA sequencing (scRNAseq) of aqueous humour cells

80-250 µl of aqueous humour was aspirated from the anterior chamber of 11 consenting patients (Supplementary Table 2) presenting at Oxford Eye Hospital with uveitis. Aqueous humour was kept at 4°C for up to 4 hours before processing for single cell RNA sequencing (scRNAseq). Cells were pelleted by centrifugation at 500x*g* for 5 mins at RT and cell pellets resuspended in 40 µl of 1% BSA/PBS and loaded onto 10X Genomics Chromium Controller (Chip K) with libraries prepared and sequenced as previously described (*44*). Gene expression and T cell receptor (TCR) sequencing libraries were prepared using the 10x Genomics Single Cell 5’ Reagent Kits v2 (Dual Index) following manufacturer’s protocol (CG000330 Rev B) and stored at -20°C at 10 nM. Libraries were sent to Novogene (Cambridge, UK) and sequenced on the NovaSeq X Plus platform (Illumina, v1.5 chemistry, 150bp paired end).

### Bioinformatic analysis of scRNAseq dataset

Sequence reads were mapped using CellRanger multi (version 7.0.0) with the 10x human reference transcriptomes (refdata-gex-GRCh38-2020-A; refdata-cellranger-vdj-GRCh38-alts-ensembl-7.1.0). Samples were merged into a single Seurat object for analysis (Seurat 4.3.01). TCR VDJ data was added to the Seurat object as metadata with scRepertoire v1.11.0. Cells with less than 200 genes or more than 10% mitochondrial content were removed. Broad cell types were predicted from raw count matrices with CellTypist using the Immune_All_High model utilising majority voting labels (*45*). Cells predicted to be T cells by CellTypist that also contained a T cell receptor were selected for T cell specific analysis. The resultant T cell specific Seurat object was scaled and samples integrated using the 2000 most variable genes and the IntegrateData() function. Dimensionality reduction was performed using the top 20 principal components. T cell subtypes were annotated from the raw T cell counts matrix with CellTypist using the Immune_All_Low model and majority voting labels (*45*). Any cell not annotated as a T cell subtype by CellTypist was relabelled as ‘other’. Diversity metrics were calculated and visualised using the clonalDiversity() function from scRepertoire. A T cell clonotype was defined as a unique combination of CDR3a and CDR3b amino acid sequences. The number of cells with identical clonotypes were calculated within each sample. Expression of tissue egress, residency and retention transcripts were compared between expanded clones (i.e., clonotypes present in more than one cell) and singlets (i.e., clonotypes only found in one cell) across each donors using the DotPlot() function in Seurat.

### Bulk RNA sequencing of trabeculectomy tissue

Iris tissue, usually discarded following surgery, was collected from donors undergoing trabeculectomy for glaucoma at either Oxford or Moorfields Eye Hospitals. The cohort consisted of 6 donors with prior history of uveitis, and 7 with no history of uveitis (Supplementary Table 4). Following surgery, tissue was placed in RPMI and kept at 4°C for up to 4 hours before being transferred to CS10 (CryoStor; 100-1061) and frozen in a Mr Frosty at -80°C and then moved to liquid nitrogen for long term storage. For RNA extraction, tissue was thawed and transferred to Buffer RLT (Qiagen) with 2-mercaptoethanol in a 2 ml soft tissue lysing kit (CK14; Precellys) and homogenised by shaking for 3 × 40 sec at 6.5 m/s on a FastPrep-24 (MP Biomedicals) with samples cooled on ice for 1 min between shakes. After homogenisation, RNA was extracted using the RNeasy Plus Micro Kit (74034; Qiagen) in accordance with the manufacturer protocol. Purified RNA was sent to Novogene and libraries were prepared using the Smart-seq2 method (*46*) and sequenced on an Illumina platform (150bp paired end).

### Weighted gene co-expression network analysis (WGCNA)

Raw reads were aligned to the human genome (GRCh38.110) and protein coding genes quantified with Rsubread (*47*). The raw counts matrix were imported into DEseq2 (*48*) R package (v 1.34.0) and normalised with the functions DESeqDataSetFromMatrix() followed by DESeq() and variance stabilisation performed with varianceStabilizingTransformation(). The 90^th^ quantile of most variable genes was selected, resulting in 2007 genes for analysis. WGCNA (*49*) was performed with the cornet pipeline (https://github.com/sansomlab/cornet) as previously described (*50*). The parameter values were set as follows: the minimum fraction of non-missing samples for a gene to be considered good was 0.5; the minimum number of non-missing samples for a gene to be considered good was 4; the minimum number of good genes was 4; the cut height for removing outlying samples was 100 (no samples removed); the minimum number of objects on a branch to be considered a cluster was 10; network type equalled signed hybrid; soft power equalled 6; adjacency correlation function was pearson; adjacency distance was calculated by euclidean distance matrix computation; topological overlap matrix type was signed; minimum module size (number of genes) was 30; and the dissimilarity threshold used for merging modules was 0.25. The analysis identified 13 distinct modules labelled with colours, plus a module of 1 unassigned gene (grey module). Module-gene membership values are specified in Supplementary Table 5. Module eigengenes (MEs), which relate to the first principal component within a given module, were calculated, and correlated with uveitis status. 5 significant (P value ≤0.05) modules were defined. Overrepresentation of GO categories and eye cell types (from the Gautam gene sets (*18*) within c8.all.v2023.1.Hs.entrez.gmt; www.gsea-msigdb.org) in module gene members were tested using 1-sided Fisher’s exact tests using gsfisher (https://github.com/sansomlab/gsfisher), comparing module gene members to the union of gene members from all the modules as the background gene set (Supplementary Table 6). Representative gene sets and pathways that had significant (BH adjusted P < 0.05) overrepresentations are displayed in the figures.

### In vivo mouse studies

#### Adoptive transfer experimental autoimmune uveitis

Adult C57BL/6J were purchased from Charles River Laboratories, UK. B6.SJL-*Ptprc*^*a*^ *Pepc*^*b*^/BoyJ (Ly5.1) mice were supplied from established breeding colonies at the University of Bristol, UK. All mice were used at 6-12 weeks of age, and confirmed negative for the Rd8 mutation (*51*), and were housed under specific pathogen free conditions with food and water *ad libitum*.

Donor Ly5.1 mice were immunised subcutaneously in both flanks with a total of 400 μg of RBP-3_629-643_ (*20*) in phosphate buffered saline (PBS) emulsified (1:1 vol/vol) with Complete Freund’s Adjuvant (CFA) further supplemented with 1.5mg/ml *Mycobacterium tuberculosis* complete H37 Ra (BD Biosciences, Oxford UK). Donors also received 1.5 μg of *Bordetella pertussis* toxin (Tocris, Bristol UK) given intraperitoneally (i.p). At 11 days post-immunisation, spleen and lymph nodes were obtained from donors and seeded at 1-2×10^6^ cells per cm^3^ in 75 cm^3^ flasks and cultured in complete medium (Dulbecco’s modified Eagle’s medium (DMEM) supplemented with 10% heat inactivated fetal calf serum (TCS Cellworks. UK), 100 U/ml penicillin-streptomycin, and 2 mmol/L L-glutamine (Invitrogen. Paisley, UK) supplemented with 10 μg/ml RBP-3_629-643_ peptide and 10 ng/ml IL-23 (R&D Abingdon, UK). 24 hours after cells are plated down the culture was further supplemented with 10 ng/ml rIL-2 (Peprotech London, UK) in complete medium. At 72 hours, leukocytes were isolated by Ficoll density centrifugation and transferred to naïve C57BL/6J by intraperitoneal injection at 2×10^6^ total leukocytes per mouse in 100 μl of PBS.

#### Clinical Imaging

Prior to imaging, pupils were dilated using topical tropicamide 1% w/v and phenylephrine 2.5% w/v (Minims; Chauvin Pharmaceuticals, Romford, UK), before anaesthesia with 2% isofluorane (Piramal Critical Care, West Drayton, UK). Posterior and anterior fundal imaging and optical coherence tomography (OCT) scans were captured using Micron IV (Phoenix Research Laboratories).

#### Dissection and dissociation

Following enucleation, single eyes were dissected in 100 µl of ice-cold PBS. Using a limbal incision the posterior segment was removed, whole retina and vitreous extracted and together with the dissecting fluid (PBS) transferred into a 1.5 ml Eppendorf tube. The tissue was mechanically dissociated by rapping the tube across an 80-well standard rack 10 times.

For anterior tissue, following lens removal, the iris, ciliary body and limbal sclera was first mechanically dissociated in a culture dish using scissors, before enzymatic digestion in 0.5 ml DMEM containing 5 mg/ml type II collagenase (Worthington, # LS004202) and 0.2 mg/ml DNase I (Roche, #11284932001) for 45 minutes at 37°C undergoing constant, gentle agitation. The enzymatic digest was stopped by adding 0.5 ml of complete media (DMEM containing 10% FCS), centrifuged at 250 *xg* for 10 mins and cell pellets resuspended in 100 µl of cold PBS.

Retina and anterior cell suspensions were transferred into a 96-well 60-µm cell filter plate (Merck Millipore) and washed with 150 µl of PBS. The plate was centrifuged at 1500 rpm for 5 min, the supernatant aspirated and cells resuspended in 0.1% bovine serum albumin (BSA) fluorescence-activated cell sorting buffer and transferred into a 96-well V-bottom plate for immunostaining.

### Flow cytometry

#### Cell surface marker staining

Cells were incubated with purified rat anti-mouse CD16/32 Fc block (1:50; 553142, [2.4G2], BD) for 10 min at 4°C before incubation with fluorochrome-conjugated monoclonal antibodies against mouse cell surface markers (Supplementary Table 10) at 4°C for 20 min. Cells were washed and resuspended in 7-aminoactinomycin D (Thermo Fisher Scientific) for dead cell exclusion.

#### Cell acquisition

Cell suspensions were acquired using a fixed and stable flow rate for 2.5 min on a four-laser Fortessa X20 flow cytometer (BD Cytometry Systems). Compensation was performed using OneComp eBeads (01-1111-41, Thermo Fisher Scientific). Seven two-fold serial dilutions of a known concentration of splenocytes were similarly acquired to construct a standard curve and calculate absolute cell numbers (*52*). Analysis was performed using FlowJo software (Treestar).

## List of Supplementary Materials

Supplementary Figure 1. tSNE plots of flow cytometry data.

Supplementary Figure 2. Cellular profiles of aqueous humour donors

Supplementary Figure 3. CD4 and CD8 T cells within phthisis bulbi retina

Supplementary Figure 4. Weighted gene co-expression network analysis (WGCNA)

Supplementary Figure 5. Disease course of adoptive transfer model of experimental autoimmune uveitis (EAU) using optical coherence tomography (OCT).

Supplementary Table 1. Details of donors used in flow cytometry experiments

Supplementary Table 2. Details of aqueous humour donors used in scRNA sequencing

Supplementary Table 3. Details of donors used in immunofluorescence imaging experiments

Supplementary Table 4. Details of trabeculectomy donors used in bulk RNA sequencing

Supplementary Table 5. WGCNA module-gene membership

Supplementary Table 6. Module gene set enrichment

Supplementary Table 7. List of antibodies used for flow cytometry on human tissue

Supplementary Table 8. Unmixed mean fluorescence intensity values of live CD3+ cells used in FlowSOM clustering

Supplementary Table 9. List of antibodies used for immunofluorescence

Supplementary Table 10. List of antibodies used for flow cytometry on mouse tissue

## Acknowledgments

We gratefully acknowledge: NHS Blood and Transplant and all patient donors and families for providing tissue to this study; Eye Research Group Oxford (ERGO); Jonathan Webber and the Kennedy Institute of Rheumatology Flow Core; Kennedy Institute of Rheumatology Digital Pathology Omics Core (DPOC). Schematic workflows were illustrated using BioRender (https://biorender.com). Dr Lindsay Nicholson for clinical scoring of mouse fundal images. We also acknowledge the flow cytometry and animal services unit facilities based at the University of Bristol.

## Author contributions

Conceptualization: ADF, IRR, CDP, SMS.

Study design: ADF, IRR, AW, LW, SNS, DAC, CDP, SMS.

Patient recruitment / sample collection: ADF, IRR, LW, TB, SAM, MA, QP, JS, SP, AB, SH, MEAAE, KB, IM, KSM, SEC, SMS.

Laboratory work: ADF, IRR, AW, LW, RH, AB, MA, DAC, SEC.

Data analysis and visualization: ADF, IRR, AW, SNS.

Interpretation: ADF, IRR, AW, LW, MA, MCC, ADD, SEC, SNS, DAC, CDB, SMS.

Funding acquisition: TB, JPS, ADD, SNS, CDP, SMS.

Project administration: ADF, CDP, SMS.

Supervision: MCC, ADD, DAC, CDP, SMS.

Writing – original draft: ADF.

Writing – review & editing: ADF, AW, LW, ADD, SNS, DAC, CPD, SMS.

## Data and materials availability

Raw fastq files used in WGCNA analysis can be accessed at http://doi.org/10.5281/zenodo.11650665. scRNAseq data can be visualized in the Broad Institute’s Single Cell Portal at https://singlecell.broadinstitute.org/single_cell/study/SCP2660. Unprocessed scRNAseq files are available upon request under a transfer agreement. All other data associated with this study are present in the paper or Supplementary Materials.

The ORBIT Consortium

In addition to ORBIT Consortium members who are authors (ADF, IRR, AW, LW, RH, JPS, ADD, SEC, DAC, CDP, SMS), the following ORBIT Consortium members are collaborators who have contributed to study design, data analysis, and interpretation:

Colin J Chu^2,4,5^

Affiliations can be found on the first page of the paper.

