## Supplementary figures for "Tissue resident memory T cells populate the human uveal tract"

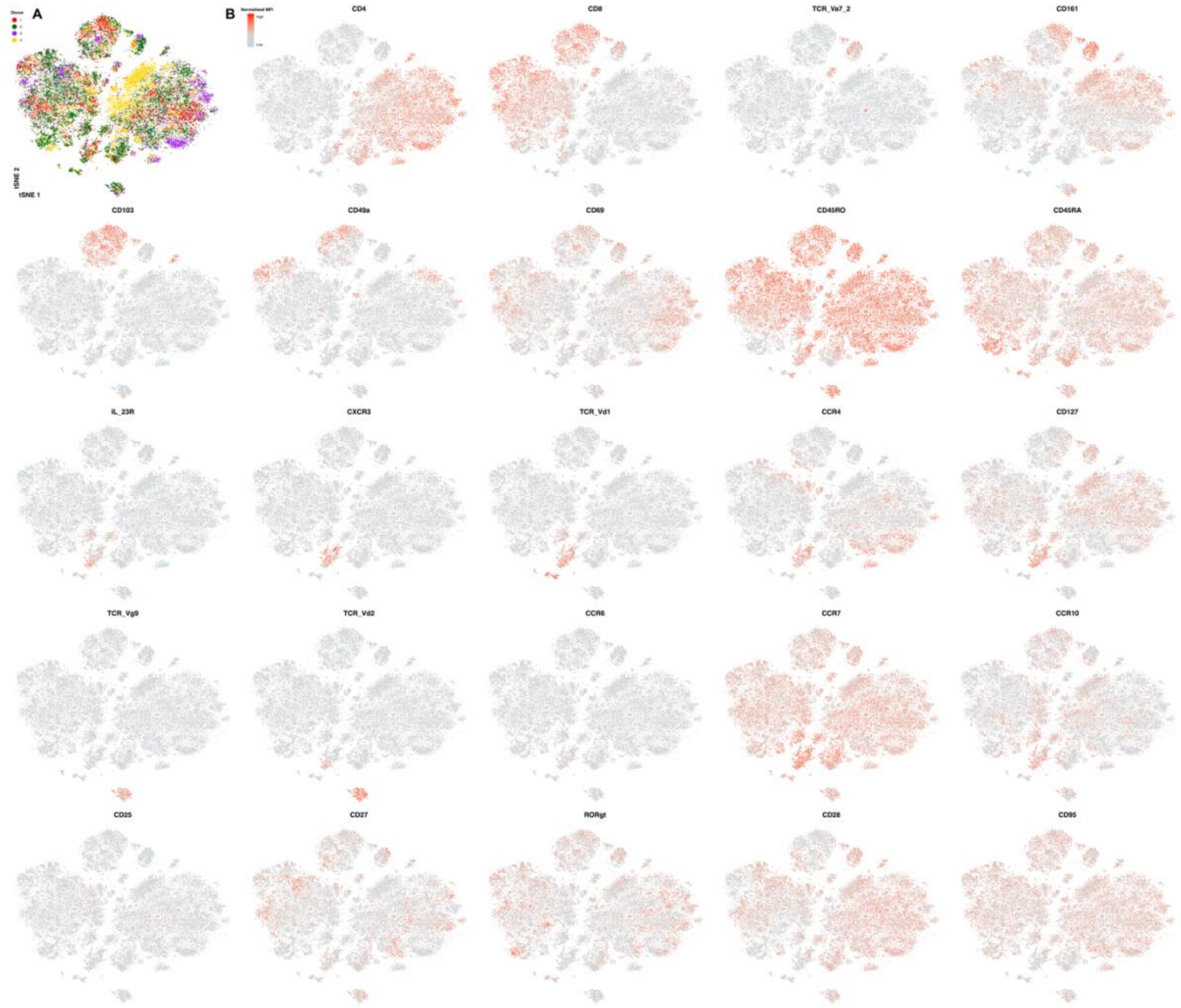


**Supplementary Figure 1. tSNE plots of flow cytometry data.**

**A**) Individual cells are coloured according to donor origin. **B**) Scaled expression for each marker gene.

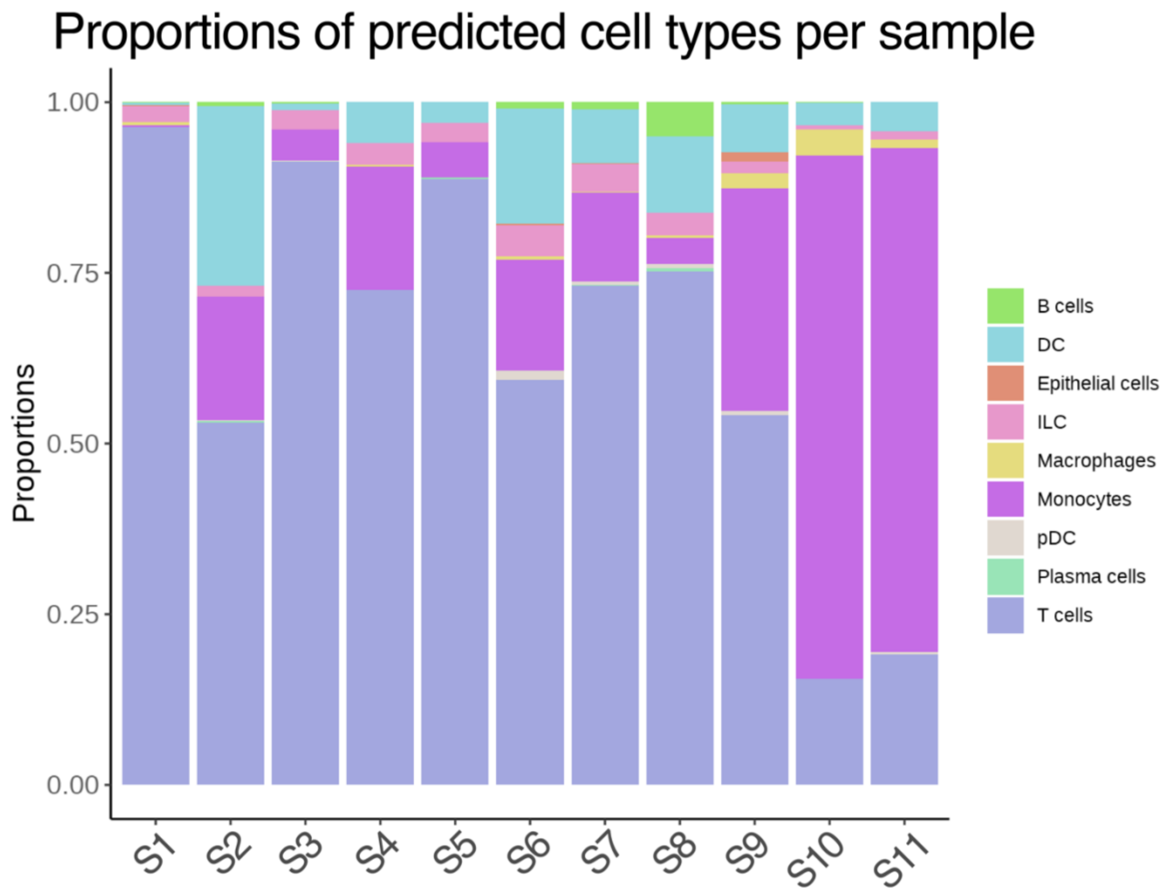


**Supplementary Figure 2. Cellular profiles of aqueous humour donors.**

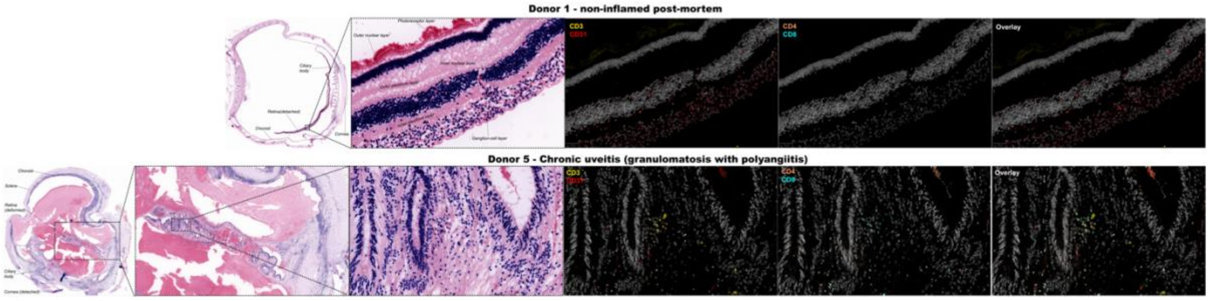


**Supplementary Figure 3. CD4 and CD8 T cells within phthisis bulbi retina**

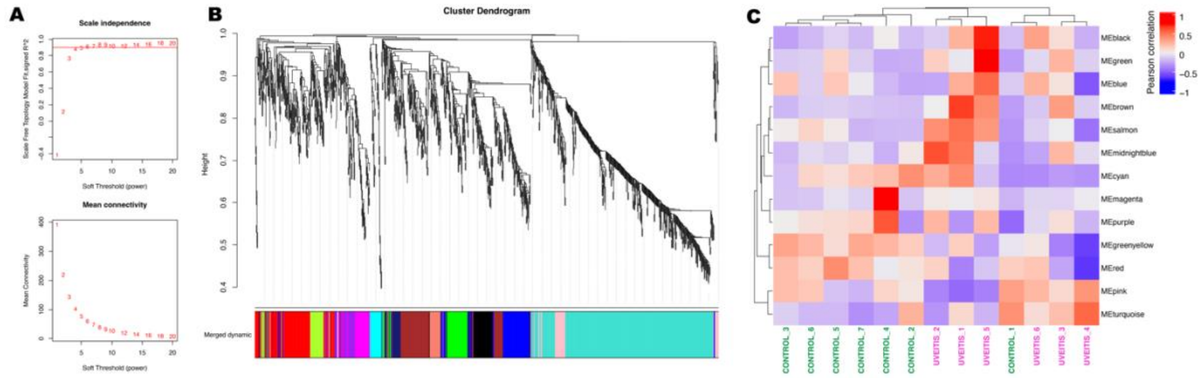


**Supplementary Figure 4. Weighted gene co-expression network analysis (WGCNA)**
**A**) Scale independence and mean connectivity plots used to select optimal soft power of 6. **B**) Dendrogram clustering genes by expression profiles with merged dynamic tree cut to select modules. **C**) Heatmap of module eigengene (ME) expression in individual donors. Donors with a history of uveitis are coloured in purple, with non-uveitic controls in green.

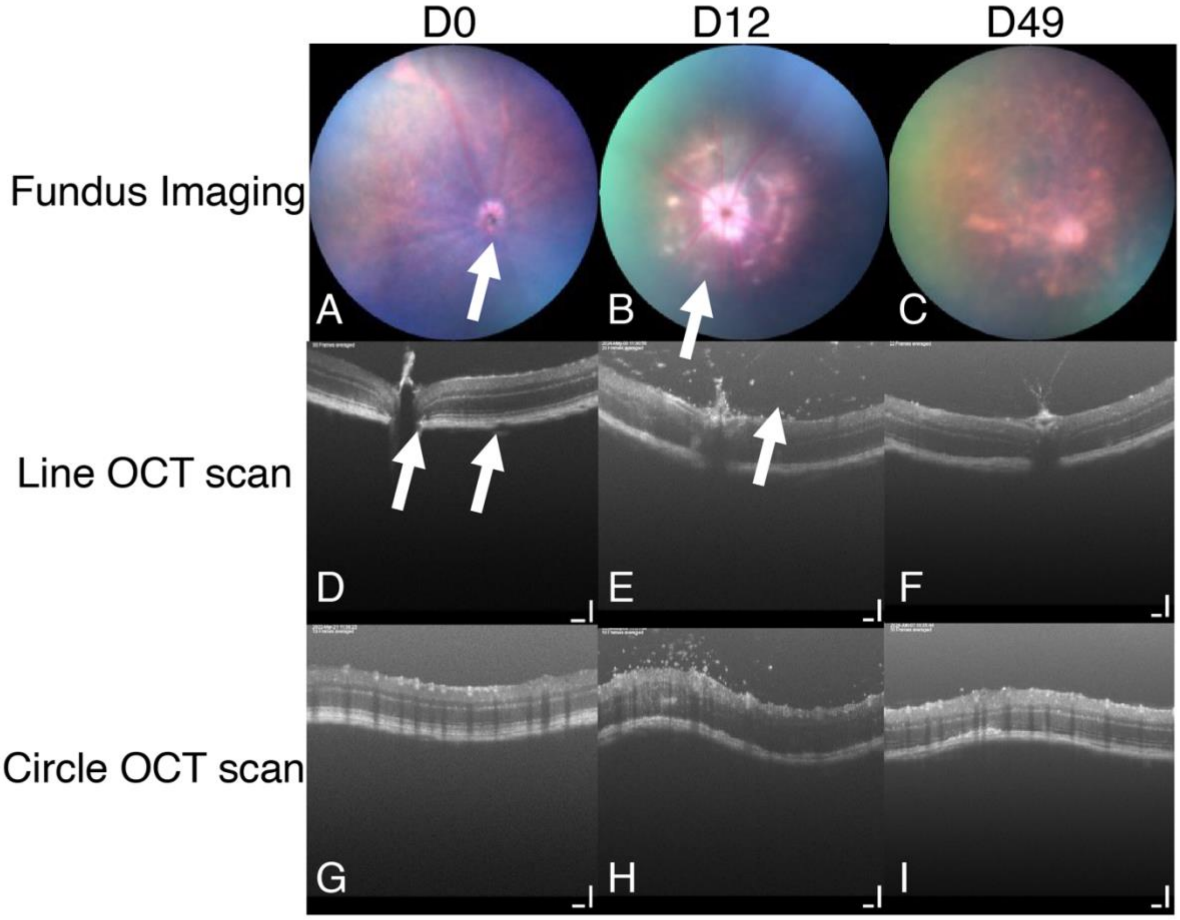


**Supplementary Figure 5. Disease course of adoptive transfer model of experimental autoimmune uveitis (EAU) using optical coherence tomography (OCT)**

**A-C)** Time course of fundus imaging throughout clinical disease with arrows to illustrate the optic disc (A) and retinal lesions (B). **D-F**) OCT line scans that follow clinical disease course, arrows illustrate optic nerve and retina (D) and infiltrating cells (E). **G-I)** OCT circle scans to match clinical disease course.
